# DWV infection and replication at the early stage *in vitro* using honey bee pupal cells

**DOI:** 10.1101/2020.09.15.297549

**Authors:** Yunfei Wu, Jing Li, Tatsuhiko Kadowaki

## Abstract

Deformed wing virus (DWV) has been best characterized among honey bee viruses; however, very little is known about the mechanisms of viral infection and replication due to the lack of honey bee cell lines. To resolve this problem, we established *in vitro* system to reconstitute DWV binding and entry to the host cell followed by translation of the genome RNA and the polyprotein processing with honey bee pupal cells. Using this system, P-domain of VP1 was found to be essential for DWV infection/replication but not binding/entry to the cell. DWV efficiently infects/replicates in cells derived from early but not late pupa, suggesting that the undifferentiated cells are targeted for the viral infection/replication. Furthermore, we found that inhibitors for mammalian picornavirus 3C-Protease, Rupintrivir and Quercetin suppress DWV infection/replication, indicating that this *in vitro* system is also useful for screening a compound to modify the viral infection/replication. Our *in vitro* system should help to understand the mechanisms of DWV infection and replication at the early stage.

**Importance:** Recent decline of managed honey bee colonies has been driven by the pathogens and parasites. However, studying the mechanisms of pathogen infection and replication in honey bee at molecular and cellular levels has been challenging. DWV is the most prevalent virus in honey bee across the globe and we established *in vitro* system to reconstitute the viral infection and replication with the primary pupal cells. Using RNA-dependent RNA polymerase (RdRP) and the negative strand of DWV genome RNA as markers, we show that the pupal cells can support DWV infection and at least replication at the early stage. The results shown in this report indicate that our *in vitro* system helps to uncover the mechanisms of DWV infection and replication. Furthermore, it is also feasible to conduct a large scale screening for compounds to inhibit or stimulate DWV infection/replication.

## Introduction

Large-scale loss of managed honey bee (*Apis mellifera*) colonies has been recently reported across the globe (Goulson et al., 2015). Since pollination by honey bees is vital for maintaining ecosystems and the production of many crops (Aizen and Harder, 2009; Klein et al., 2007), prevention of honey bee colony losses has become a major focus in both apiculture and agriculture. Colony losses have often been associated with the ectoparasitic mites *Varroa destructor* and *Tropilaelaps mercedesae*, which feed on honey bees and transmit honey bee viruses, particularly deformed wing virus (DWV) to the host (Chantawannakul et al., 2018; de Miranda and Genersch, 2010; Rosenkranz et al., 2010). In the absence of mites, DWV copy numbers remain low in honey bees without specific symptoms (covert infection). However, DWV levels associated with honey bees are dramatically increased in the mite infested colonies (Forsgren et al., 2009; Khongphinitbunjong et al., 2016; Shen et al., 2005; Wu et al., 2017). These honey bees often show multiple symptoms (overt infection), which include the death of pupae, deformed wings, shortened abdomen, and reduced lifespan (de Miranda and Genersch, 2010; Rosenkranz et al., 2010; Tentcheva et al., 2004; Yue et al., 2007). Thus, winter colony loss is strongly correlated with the presence of DWV and *V. destructor* (Highfield et al., 2009; Nazzi and Le Conte, 2016).

DWV belongs to the Iflaviridae family and exists as a nonenveloped icosahedral virion about 30 nm in diameter, which contains a positive-strand RNA genome of ∼10,000 nt. The genome RNA is translated to a polyprotein which is co-translationally and post-translationally cleaved by the viral protease to produce structural and nonstructural proteins (Lanzi et al., 2006). DWV virion is constructed from the VP1, VP2, and VP3 which are arranged into a capsid with a pseudo-T3 icosahedral symmetry. The C-terminus of VP1 (P-domain) is present at the outermost of virion and undergoes conformational change under different pH condition, suggesting that it may act as catalytic site to enable viral entry into the host cell (Organtini et al., 2017; Skubnik et al., 2017).

Although DWV has been best characterized among honey bee viruses, very little is known about how the virus binds, enters, and replicates in the host cell. DWV can propagate in honey bee larva and pupa by the viral injection (Gusachenko et al., 2020; Lamp et al., 2016; Ryabov et al., 2020); however, this *in vivo* system does not allow us to study the underlying mechanisms of viral infection and replication. Honey bee cell line would provide the best resource to study virus and other pathogens (Guo et al., 2020) but it has not been available to date. To solve this problem, we developed *in vitro* system to reconstitute DWV infection and replication at the early stage using primary cells derived from honey bee pupa. The mechanistic insight into DWV infection and replication obtained by our *in vitro* system will be reported and discussed in this study.

## Results

### DWV infection and replication in honey bee pupal cells

To establish a method to characterize DWV infection and replication in honey bee cell *in vitro*, we dissected head and abdomen of pupa with pale or pink eyes and infected the halves of tissues with purified DWV in the culture medium. We then tested the degree of viral infection and replication by quantifying one of the non-structural proteins, RNA-dependent RNA polymerase (RdRP). As shown in Figure 1A, two bands corresponding to RdRP precursor with 3C-protease (3C-Pro) (90 kDa) and matured RdRP (53 kDa) were specifically detected in the head and abdominal tissues infected by DWV. RdRP precursor was more abundant than the matured protein, suggesting that the cleavage between RdRP and 3C-Pro is rate-limiting as other picornaviruses (Jiang et al., 2014). RdRP precursor was not detected at 6 h but increased at 12 and 24 h after infection of the pupal head cells (Fig. 1B and C). Furthermore, it was possible to detect the negative strand RNA of DWV genome (Fig. 1D), and thus the viral replication was also initiated. These results demonstrate that DWV infects and starts replicating in honey bee pupal cells *in vitro*. Using RdRP as a marker, we could study the mechanisms of viral infection and replication at early stage.

**Figure 1.**
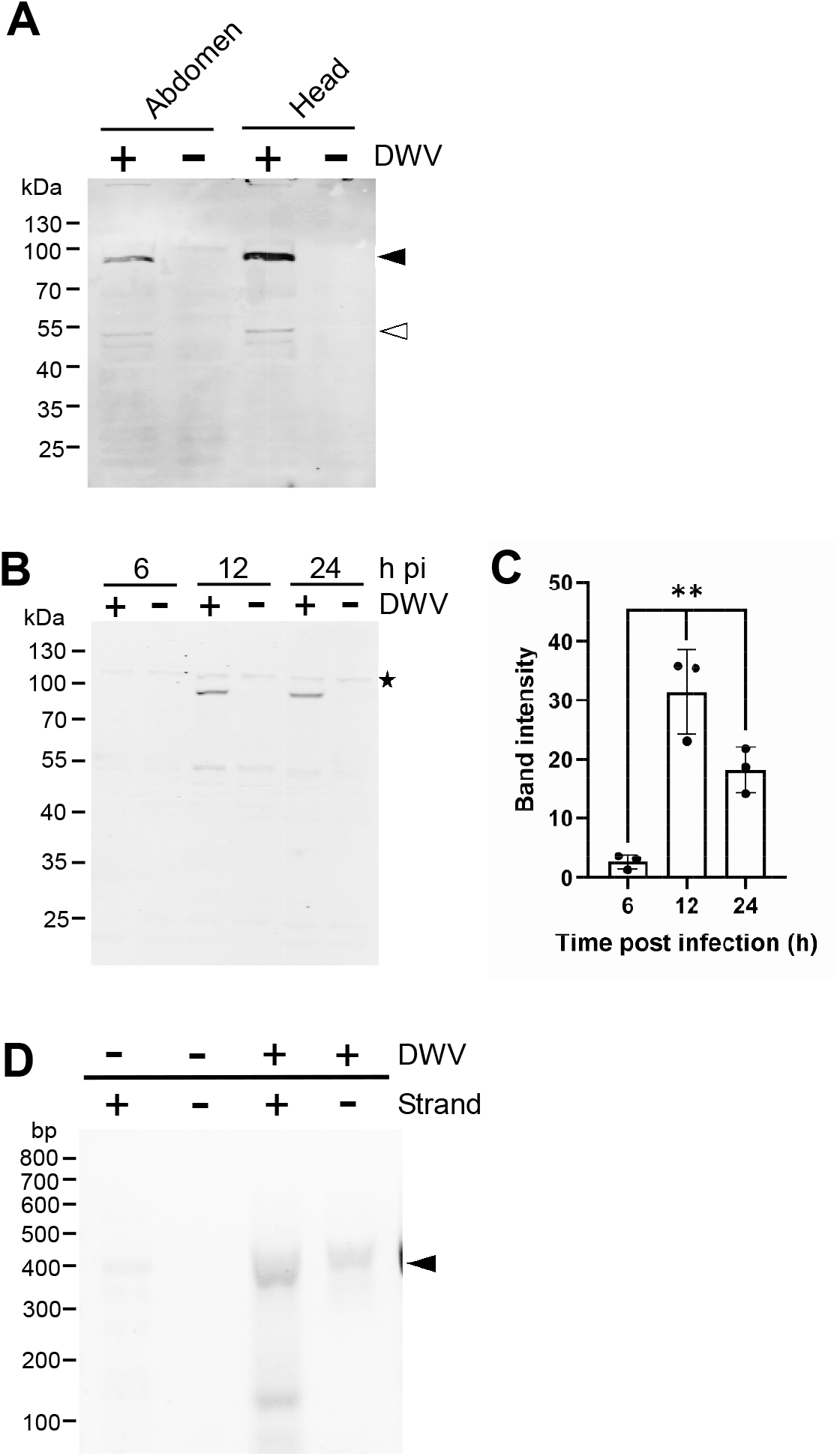
DWV infection and replication in honey bee pupal cells. (A) Abdomen and head from the single pupa were dissected to half and one was infected by DWV (+) and the other was left untreated (-). DWV infection/replication was tested by western blot using anti-RdRP antibody. RdRP precursor with 3C-Protease (90 kDa) and mature RdRP (53 kDa) bands are indicated by black and white arrowheads, respectively. The size (kDa) of protein molecular weight marker is at the left. (B) The synthesis of RdRP in pupal head cells was tested at 6, 12, and 24 h after DWV infection (h pi). The star represents non-specific protein cross reacted with anti-RdRP antibody. (C) Band intensity of RdRP precursor is compared between three time points by Dunnett test (one-tailed) and *P*-values between 6 and 12 or 24 h are < 0.0003 and < 0.007, respectively (_**_). (D) Positive (+) and negative (-) strands of DWV genome RNA in pupal head cells with (+) or without (-) DWV infection were detected by the strand-specific RT-PCR. PCR amplicon with the expected size is indicated by black arrowhead. The size (bp) of DNA molecular weight marker is at the left.

### DWV infection and replication in honey bee pupal cells at the different developmental stages

We next tested whether DWV infection and replication in honey bee pupal cells depends on the developmental stage. We infected the head cells from pupae with pale eyes, purple eyes, yellow thorax, and brown thorax by DWV. RdRP precursor decreased with the cells from the pupae at later developmental stages (Fig. 2A and B). These results demonstrate that DWV infection and/or replication becomes inefficient in honey bee head cells by the progression of pupal development.

**Figure 2.**
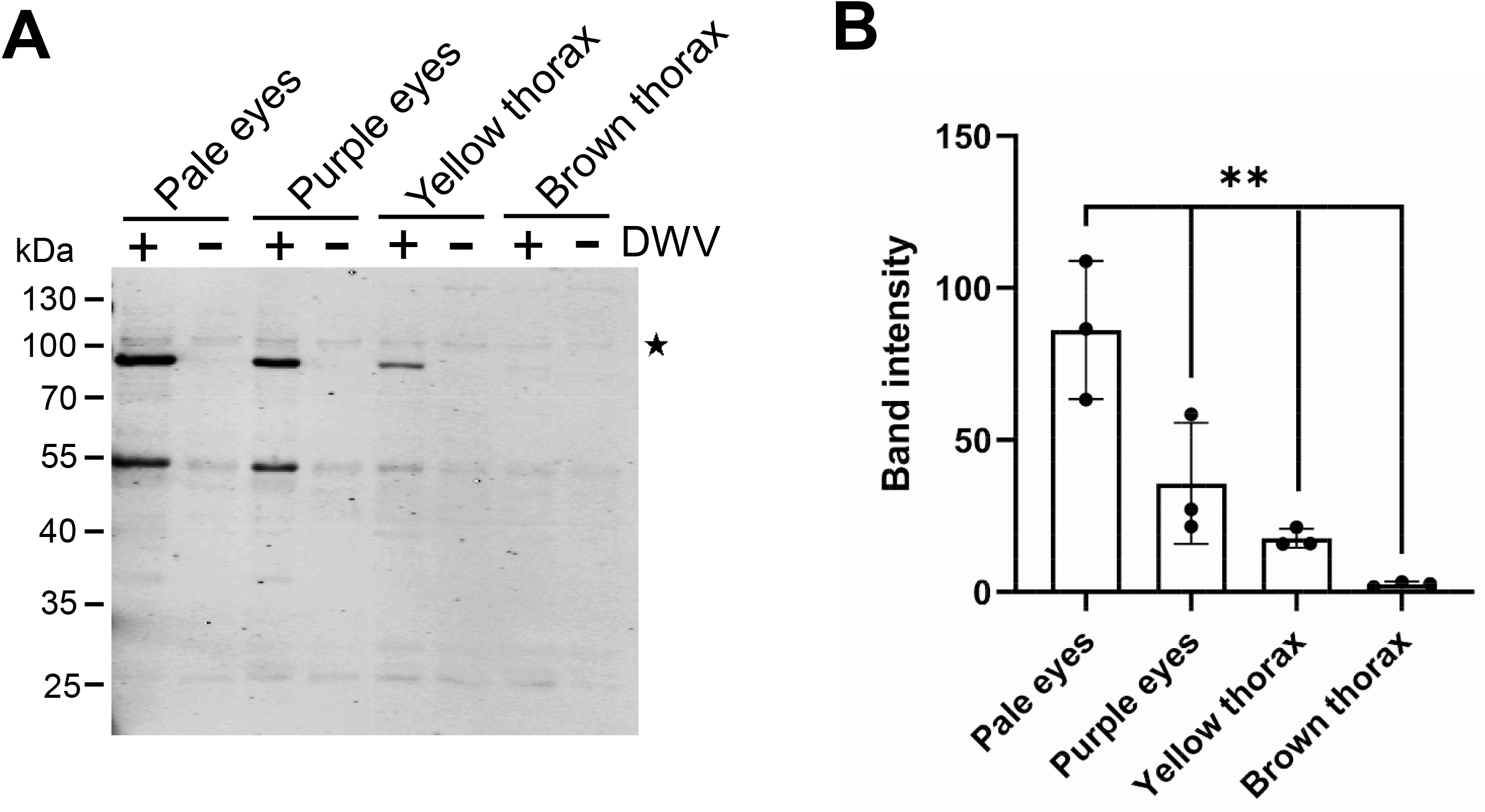
DWV infection and replication in honey bee pupal cells at the different developmental stages. (A) Pupal head cells at four different developmental stages (pupa with pale eyes, purple eyes, yellow thorax, and brown thorax) were infected by DWV, and then RdRP synthesis was tested. (B) Band intensity of RdRP precursor is compared between the four groups by Dunnett test (one-tailed) and *P*-values between pale eyes and purple eyes, yellow thorax, or brown thorax are < 0.005, < 0.0008 and < 0.0002, respectively (_**_).

### Roles of VP1 P-domain for DWV infection/replication

The structural analysis of DWV virion by X-ray crystallography and Cryo-EM showed that P-domain of VP1 (amino acid 748-901 of DWV polyprotein) was present at the outermost surface of virion and suggested to bind the viral receptor or disrupt membrane to deliver its genome RNA into cytosol (Organtini et al., 2017; Skubnik et al., 2017). To understand the roles of P-domain for DWV infection/replication using our *in vitro* system, we pre-incubated DWV with anti-VP1 P-domain antibody and then infected the pupal head cells. This antibody binds DWV virion under native condition because it immunoprecipitated VP1, VP2, and VP3 (Supplementary Figure). Another antibody recognizing the different domain of VP1 (amino acid 524-750 of DWV polyprotein) which does not bind DWV virion under native condition was used as a control. Increasing amount of anti-VP1 P-domain but not anti-VP1 (524-750) antibody suppressed the synthesis of RdRP precursor in the pupal head cells (Fig. 3A and B). These results indicate that P-domain of VP1 plays essential roles for DWV infection and/or replication.

**Figure 3.**
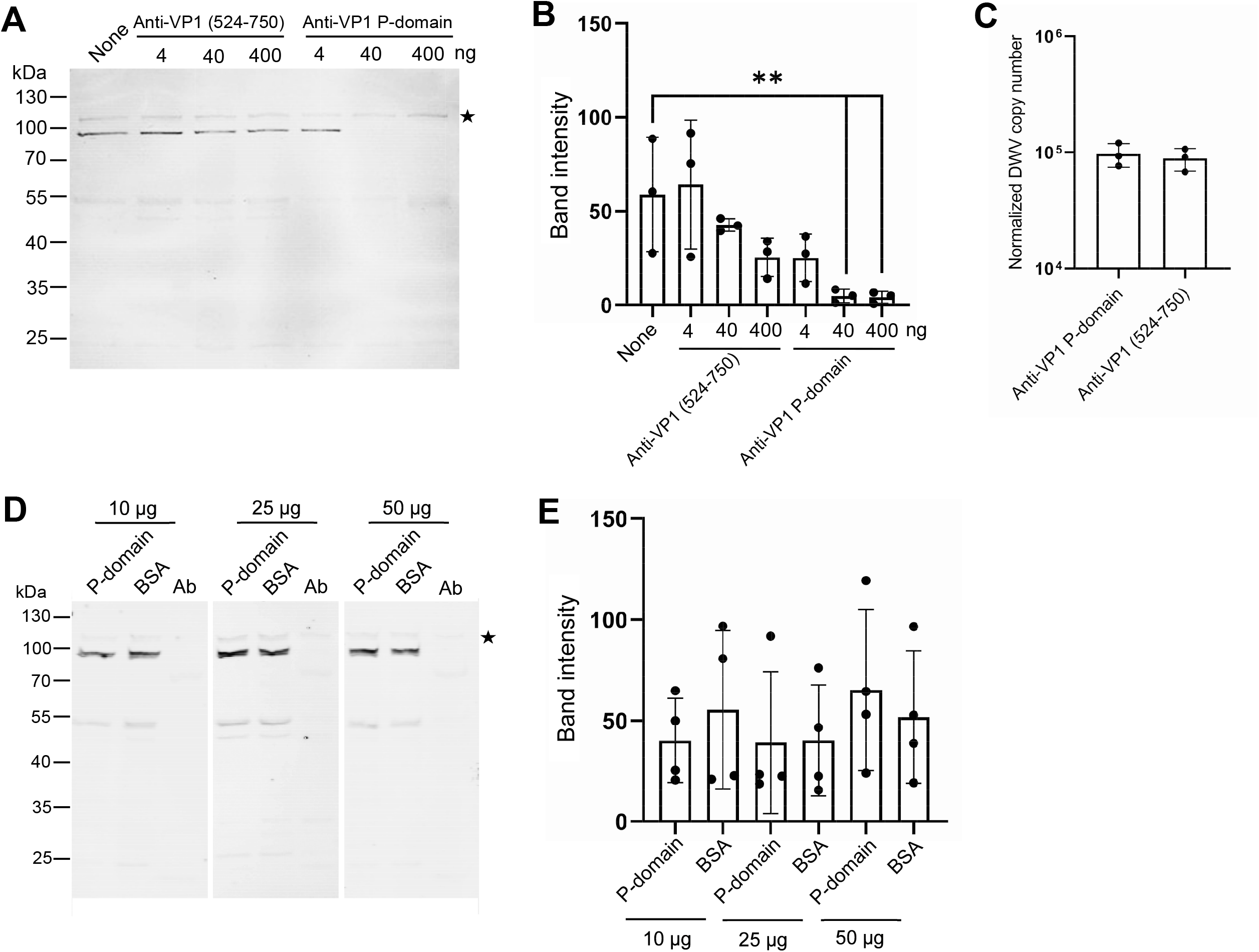
Essential roles of VP1 P-domain for DWV infection/replication. (A) The effects of pre-incubating DWV with 4, 40, or 400 ng of either Anti-VP1 (524-750) or Anti-VP1 P-domain antibody before infection on RdRP synthesis were tested. DWV without the antibody pre-incubation (None) was used as a control. (B) Band intensity of RdRP precursor is compared between the seven groups by Dunnett test (one-tailed) and *P*-values between None and 40 or 400 ng of Anti-VP1 P-domain antibody are < 0.008 and < 0.007, respectively (_**_). (C) Entry of DWV pre-incubated with 40 ng of either Anti-VP1 (524-750) or Anti-VP1 P-domain antibody to pupal head cells (Copy number of DWV inside the infected cells) was compared. There is no statistically significant difference between the two groups. (D) The effects of pre-incubating pupal head cells with 10, 25, or 50 μg of purified P-domain protein before DWV infection on RdRP synthesis were tested. BSA was used as a control. Lysate of abdomen from the same pupa (Ab) was also analyzed to confirm the lack of replication of endogenous DWV. (E) Band intensity of RdRP precursor is compared between P-domain and BSA with the different amount of protein. There is no statistically significant difference between the two groups.

To uncover the mechanism of inhibiting DWV infection/replication by anti-VP1 P-domain antibody, we tested entry of DWV preincubated by the antibody to pupal head cells. As shown in Figure 3C, DWV entry to the cells was comparable between treatments by anti-VP1 P-domain and anti-VP1 (524-750) antibodies, indicating that masking P-domain with the antibody does not affect the viral binding and entry. Furthermore, we also found that preincubating pupal head cells with purified P-domain protein prior to DWV infection does not affect RdRP precursor synthesis (Fig. 3D and E). These results demonstrate that P-domain of VP1 is essential for DWV infection/replication but not binding the viral receptor or the following entry to cell.

### Identification of inhibitors for DWV infection/replication

Our *in vitro* system allowed us to identify inhibitors for DWV infection and replication at the early stage. We thus tested the effects of known inhibitors for mammalian picornavirus (Baggen et al., 2018; Owino and Chu, 2019) on DWV infection/replication. As shown in Figure 4A and B, 3C-Pro inhibitors, Rupintrivir (Binford et al., 2005) and Quercetin (Yao et al., 2018) decreased the synthesis of RdRP precursor at 5 μM and 5 mM, respectively. This is consistent with the essential roles of 3C-Pro for cleavage of the viral polyprotein and host proteins (Sun et al., 2016). Both compounds did not dramatically affect the viability of pupal head cells at these concentrations (Fig. 4C), suggesting that they suppressed the synthesis of RdRP precursor by specifically inhibiting 3C-Pro. These results show our *in vitro* system using honey bee pupal cells can be used to identify inhibitors as well as stimulators for DWV infection/replication.

**Figure 4.**
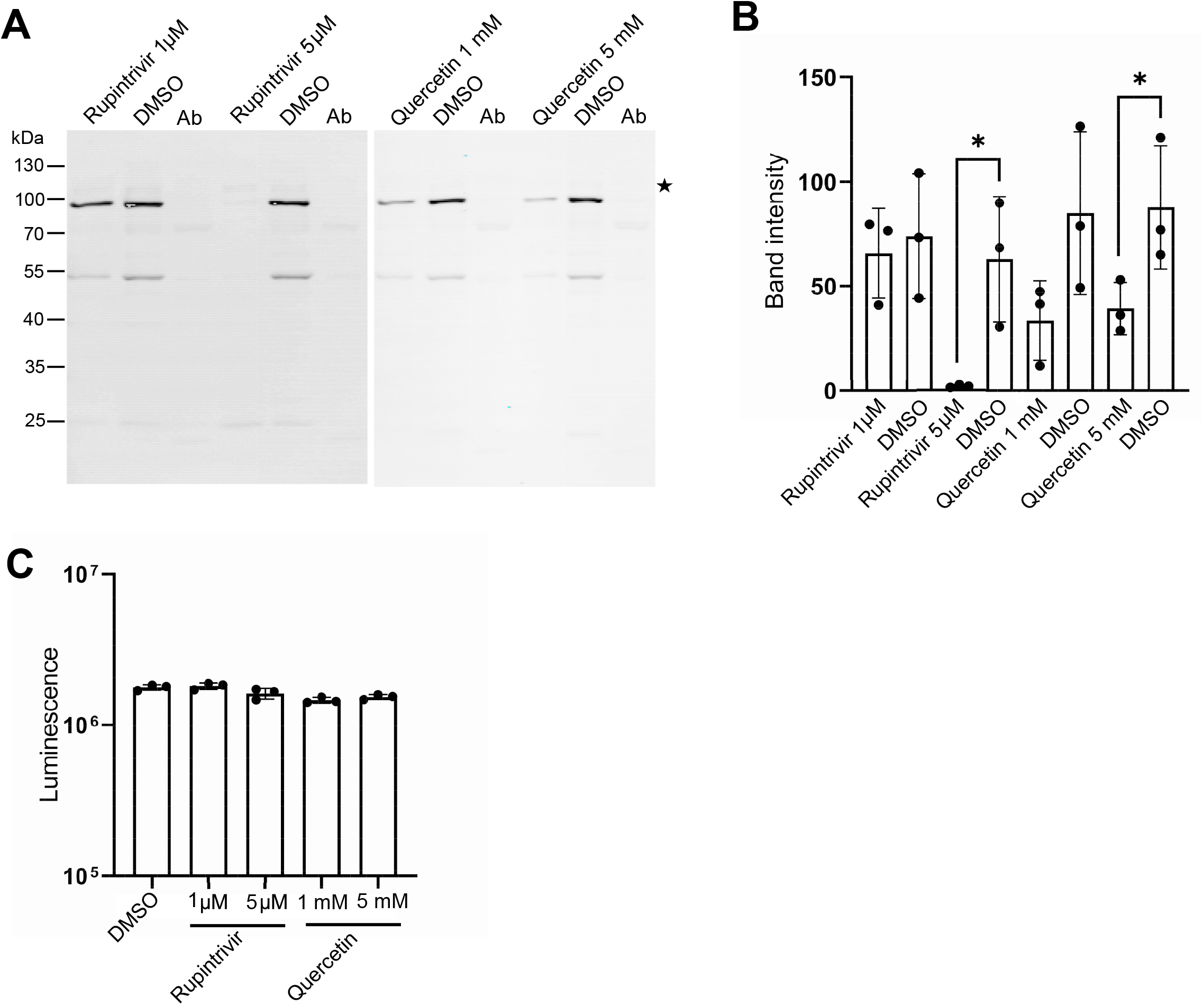
Rupintrivir and Quercetin suppress RdRP synthesis in DWV-infected cells. (A) The effects of Rupintrivir (1 and 5 μM) and Quercetin (1 and 5 mM) on RdRP synthesis in DWV-infected pupal head cells were tested. DMSO was used as a control. (B) Band intensity of RdRP precursor is compared between DMSO and Rupintrivir or Quercetin at the different concentration by Welch’s *t*-test (one-tailed) and *P*-values between DMSO and 5 μM Rupintrivir or 5 mM Quercetin are < 0.04 and < 0.05, respectively (_*_). (C) Luminescence generated by luciferase activity dependent on the intracellular ATP level is compared between the pupal head cells treated with DMSO, Rupintrivir or Quercetin at the indicated concentrations.

## Discussion

Using RdRP as a marker, we demonstrated that purified DWV infects and starts replicating in the cultured honey bee pupal cells. We also confirmed the synthesis of negative strand RNA of DWV genome but not new virions in the infected cells. The primary pupal cells are unlikely to support the long-term viral replication. Nevertheless, using this *in vitro* system, it is possible to study the mechanisms of DWV infection and replication at early stage: 1) binding and entry of DWV to honey bee cell; 2) uncoating the viral genome RNA to cytosol; 3) translation of viral genome RNA followed by the polyprotein processing.

DWV infection/replication becomes inefficient with pupal head cells at the late development. These results suggest that DWV efficiently infects and replicates in the undifferentiated but not differentiated cells. The host factors necessary for the viral infection/replication may decrease upon cell differentiation. This is consistent with the previous study reporting that higher dose of injection is required for adult bees than pupae to detect DWV later (Mockel et al., 2011). Thus, DWV efficiently infects/replicates in early pupa when introduced by the parasitic mite (both *V. destructor* and *T. mercedesae*) during the reproductive phase but not in adult by *V. destructor* during the phoretic phase. Introducing DWV in early pupa must be a critical factor for the increase of viral load by the mite infestation. This property may explain why DWV and perhaps other honey bee viruses are generally benign and often establish the persistent (covert) infection. DWV appears to actively infect and replicate only in early embryo and pupa containing many undifferentiated stem cells. However, once these cells differentiate to the specific cell types, DWV remains associated with the cells without active infection and replication in other cells. DWV is present in various cells and tissues throughout the whole body of honey bee (Mazzei et al., 2014; Mockel et al., 2011; Shah et al., 2009; Zioni et al., 2011) and they are likely to be the progenies of stem cells infected by DWV. DWV is usually present as a covert swarm of the variants; however, mite infestation was shown to dramatically decreases the viral diversity (Martin et al., 2012). This could happen by the selective replication of DWV variant exogenously introduced by the mite followed by taking over the viral population in early pupa.

We showed that masking P-domain of VP1 with the antibody inhibits DWV infection/replication without affecting the entry to honey bee cell. These results suggest that P-domain does not appear to bind the viral receptor on the cell surface. This is also consistent with the lack of inhibiting DWV infection/replication by preincubation of pupal cells by the purified P-domain protein. P-domain is likely to be necessary for the release of DWV genome RNA to cytosol. Consistent with this hypothesis, P-domain was suggested to contain the putative catalytic amino acids for lipase, protease as well as esterase and undergo large conformational movement under low pH condition (Organtini et al., 2017; Skubnik et al., 2017). DWV may bind honey bee cell nonspecifically followed by the entry through endocytosis, and then P-domain creates a pore in the endosomal membrane to deliver the genome RNA to cytosol for the translation.

Using our *in vitro* system, we demonstrated that 3C-Pro inhibitors for mammalian picornavirus, Rupintrivir and Quercetin, reduce the synthesis of RdRP precursor and thus suppress DWV replication. Both compounds were shown to directly bind 3C-Pros of rhinovirus and enterovirus (Costenaro et al., 2011; Lu et al., 2011; Wang et al., 2011; Yao et al., 2018), suggesting that DWV 3C-Pro shares the similar structure as well as the amino acids interacting with the compounds. Nevertheless, Rupintrivir and Quercetin are effective at much lower concentrations (in range of nM and μM, respectively) against infection of mammalian picornaviruses (Binford et al., 2005; Yao et al., 2018) than DWV. Once structure of DWV 3C-Pro becomes available, it would be possible to design more potent inhibitors by modifying above compounds. We also tested the effects of other inhibitors, for example, Arbidol (Herod et al., 2019) and 6,7-dichloroquiloline-5,8-dione (Jung et al., 2018); however, they did not affect the synthesis of RdRP precursor. These results indicate that our *in vitro* system will be useful for screening compounds to affect DWV infection and replication at early stage. It should also help to uncover the mechanism of DWV infection/replication in honey bee cell. It is, nevertheless, still useful to establish honey bee cell lines to reproduce the complete cycle of DWV infection and replication *in vitro*.

## Materials and Methods

### Purification of DWV

46 pale-eyes pupae from a mite-free colony were injected with DWV (10^5^ copy/pupa) and incubated at 33 °C for 2 days. They were homogenized with 50 mL PBS, and then the supernatant was collected after centrifugation. The supernatant was filtered by 0.22 μm nylon membrane followed by centrifugation through a cushion of 50 % (w/v) sucrose with 36,000 rpm at 4 °C for 4 h (Beckman SW41 Ti rotor). The pellet was resuspended with PBS and the copy number of DWV was quantified by qRT-PCR.

### Quantification of DWV by qRT-PCR

DWV copy number was determined by qRT-PCR using a Hieff^™^® qRT-PCR SYBR Green Master Mix (Low Rox Plus, Yesen) and DWV #1 primers (Supplementary Table). To prepare a standard curve for DWV, PCR product obtained by above primers was purified and the copy number was determined by a formula below.

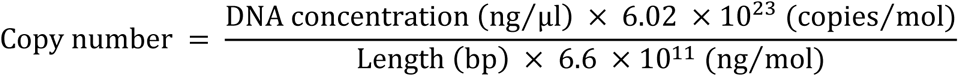

6.6 × 10^11^ ng/mol is the average molecular mass of one base pair and 6.02 × 10^23^ copies/mol is Avogadro’s number. We conducted qPCR using 10^1^-10^9^ copy number of the PCR product and then plotted the Ct values against the log values of copy numbers. DWV copy number in the sample was determined using the standard curve. The amount of cDNA added to each qPCR reaction was normalized using *A. mellifera 18S rRNA* as the endogenous reference (Supplementary Table).

### Raising anti-VP1 (524-750) antibody

The partial VP1 cDNA corresponding to amino acid 524-750 of DWV polyprotein was amplified by PCR using the primer sets, 5’-SacI-VP1 and 3’-Hind?-VP1 (Supplementary Table). The amplified PCR product was digested by SacI (NEB) and Hind? (NEB), and then subcloned to pCold I vector (TAKARA) followed by transformation to BL21. The transformed BL21 was grown in 1L of LB medium containing 1 % glucose and 0.1 mg/mL Ampillicin at 37°C until A_600_ reached to 0.5. The cell suspension was cooled down, and then IPTG was added at 0.5 mM to induce the protein expression at 15 °C for 16 h. *E. coli* was collected by centrifugation and resuspended in 100 mL of 50 mM NaH_2_PO_4_, 300 mM NaCl, 1 mM DTT, 0.1 % sarkosyl, pH 8.0 containing protease inhibitors. The cell lysate was prepared by sonication using Q700 Sonicator (Qsonica) at amplitude 100 on ice for 45 min (30 sec pulse with 3 min-interval). The supernatant was collected by centrifugation and His-tag protein purification resin (Beyotime) was then added to the supernatant. After gently rotating at 4 °C for 2 h, the resin was washed five times with 10 mL of 50 mM NaH_2_PO_4_, 300 mM NaCl, 2 mM imidazole, 0.1 % sarkosyl, pH 8.0. The bound protein was eluted with 16 mL of 50 mM NaH_2_PO_4_, 500 mM NaCl, 250 mM imidazole, 0.1 % sarkosyl, pH 8.0. The eluted protein was dialyzed against 2L of PBS with 0.1 % sarkosyl, and then concentrated using Vivaspin^®^ 6 polyethersulfone 10 kDa (Sartorius). The purified and concentrated protein was delivered to GeneScript-Nanjing to raise the anti-rabbit polyclonal antibody.

### Infection of honey bee pupal cells by DWV

Honey bee pupae with pale/pink-eyes were first collected from a mite-free colony if not indicated. They were surface sterilized by washing with 10 % bleach followed by sterile PBS three times (5 min for each wash). The head was dissected to approximately half and each piece was suspended in 100 μL of Grace medium containing 10 % FBS and antibiotics (penicillin and streptomycin) with or without DWV (3 × 10^4^ copy) at 33 °C for 1 h. Fresh culture medium (400 μL) was then added in 24-well plate and incubated at 33 °C for 16h if not indicated. The pupal head cells were collected and washed three times with PBS followed by homogenization with 150 μL RIPA buffer (20 mM Tris-HCl, pH 7.5, 150 mM NaCl, 1 % NP-40, 0.5 % sodium deoxycholate, 0.1 % SDS) containing protease inhibitors. The pupal abdominal cells were homogenised with 400 μL of above buffer. After centrifugation, the protein concentration of supernatant was measured using Enhanced BCA protein assay kit (Beyotime). The cell lysates with the equal amount of protein were analyzed by western blot. The abdominal cells were simultaneously tested to confirm the lack of replication of endogenous DWV in the pupa.

DWV was pre-incubated with either anti-VP1 P-domain (Wu et al., 2020) or anti-VP1 (524-750) antibody at the indicated amount of protein for 30 min, and then added to the pupal head cells for infection as above. The dissected pupal head cells were pre-incubated with either purified VP1 P-domain protein (Wu et al., 2020) or BSA at the indicated amount of protein in the Grace culture medium for 1 h. DWV was then added for infection as above. Pre-incubation was conducted at room temperature. Rupintrivir, Quercetin, or DMSO was added to the pupal head cells at the indicated concentration together with DWV as above.

### Western blot

The protein samples in SDS-PAGE sample buffer (2 % SDS, 10 % glycerol, 10 % β-mercaptoethanol, 0.25 % bromophenol blue, 50 mM Tris-HCl, pH 6.8) were heated at 99 °C for 5 min. After centrifugation, the supernatants were applied to 10 % SDS-PAGE and the proteins were transferred to a nitrocellulose membrane (Pall^®^ Life Sciences). The membrane was then blocked with PBST (PBS with 0.1 % Tween-20) containing 5 % BSA for 1 h at room temperature followed by incubating with 1000-fold diluted anti-RdRP antibody (Wu et al., 2020) at 4 °C overnight. The membrane was washed three times with PBST (5 min each), and then incubated with 10,000-fold diluted IRDye® 680RD donkey anti-rabbit IgG (H+L) (LI-COR Biosciences) in PBST containing 5 % skim milk at room temperature for 1.5 h. The membrane was washed as above, and then visualized using Odyssey Imaging System (LI-COR Biosciences). Band intensity of 90 kDa RdRP precursor was measured by image-J.

### Isolation of total RNA and RT-PCR

Total RNA was extracted from the cultured pupal head cells using TRI Reagent® (Sigma-Aldrich) according to the manufacturer’s instruction. To detect DWV genome RNA, reverse transcription (RT) reaction was carried out using 1 μL of total RNA, random primer (TOYOBO), ReverTra Ace (TOYOBO), and RNase inhibitor (Beyotime). RNase H (Beyotime) was then added to digest RNA in RNA/cDNA heteroduplex after cDNA synthesis. PCR was conducted using F15 and B23 primers (Supplementary Table) and the cycling condition of 2 min at 94 °C followed by 30 cycles of 10 sec at 98 °C, 20 sec at 55 °C, and 30 sec at 68 °C. To detect the negative strand of DWV genome RNA, RT was carried out as above except Tag-F15 primer (Supplementary Table) was used instead of random primer. PCR was performed using Tag and B23 primers (Supplementary Table) as above. The PCR products were analysed by 2 % agarose gel.

### Quantification of DWV entered to the cells

DWV was pre-incubated with either 40 ng of anti-VP1 P-domain or anti-VP1 (524-750) antibody for 30 min, and then added to the pupal head cells in the Grace culture medium at 33 °C for 2 h. The cells were then washed three times with PBS followed by treating with 0.5 % trypsin at room temperature for 10 min to remove DWV on the cell surface. After washing three times with PBS, total RNA was extracted and DWV genome RNA was quantified by qRT-PCR as above.

### Testing cell viability

Heads from ten pale-eyes pupae were dissected and homogenized 15 times with 10 mL Grace culture medium using Dounce homogenizer (Loose fitting). The homogenate was then filtered through a cell strainer (Falcon) and 100 μL was inoculated to each well in 96-well plate. The cells were cultured in the presence of either DMSO, Rupintrivir (1 and 5 μM), or Quercetin (1 and 5 mM) at 33 °C for 16 h. The cultured plate was incubated at room temperature for 10 min followed by adding 100 μL of reagent solution (CellTiter-Lumi™ Luminescent Cell Viability Assay Kit, Beyotime). The plate was shaken at room temperature for 2 min to promote cell lysis, and then further incubated for 10 min to stabilize the Chemiluminescence signal. The signal was detected using Varioskan™ LUX multimode microplate reader (Thermo Fisher) and depends on intracellular ATP level, thus the relative viability of cells.

### Mass spectrometry analysis of immunoprecipitates by anti-VP1 P-domain antibody

Anti-VP1 P-domain antibody (75.4 μg) or pre-immune serum was first bound to Protein A-agarose (Beyotime), and then washed three times with 0.2 M sodium borate (pH 9.0). The cross-linking was conducted by mixing 20 mM Dimethylpimelimidate (Thermo Fisher) thoroughly and incubating at room temperature for 40 min with agitation. To quench the reaction, 0.2 M ethanolamine (pH 8.0) was added and agitated for 1 h. The cross-linked beads were washed three times with 150 mM NaCl containing 0.58 % acetic acid followed by three washes with ice-cold PBS. The lysates of abdomen from six pale-eyes pupae injected by DWV were prepared using 2 mL RIPA buffer containing protease inhibitors The beads were incubated with the above cell lysates at 4 °C overnight, and then washed three times with buffer containing 150 mM NaCl, 50 mM Tris-HCl, pH 8.0, 10 mM EGTA, 0.2 % NP-40 and additional three washes with 50 mM ammonium bicarbonate. The protein-bound beads in 50 μL of 50 mM ammonium bicarbonate were heated at 95 °C for 5 min. The immunoprecipitates were applied to 8 % SDS-PAGE and the gel was stained by ProteoSilver™ stain kit (Sigma-Aldrich) according to manufacturer’s instruction. Four major bands specifically immunoprecipitated by anti-VP1 P-domain antibody were cut into small pieces and sequentially destained by 100 mM ammonium bicarbonate, 100 mM ammonium bicarbonate/acetonitrile (1:1 mixture), and 100 % acetonitrile. The gel pieces were then reduced by 10 mM DTT in 100 mM ammonium bicarbonate solution for 1 h at 56 °C and alkylated by 55 mM iodoacetamide in 100 mM ammonium bicarbonate solution for 45 min at room temperature under dark. They were dried then swelled in 25 mM ammonium bicarbonate containing 20 ng trypsin on ice followed by overnight incubation at 37 °C. Mass spectrometric detection was performed using Easy-nLC 1,000 coupled to LTQ Orbitrap Elite mass spectrometer (Thermo Fisher). It was equipped with nanoelectrospray source and operated in data-dependent acquisition (DDA) mode with the following settings: spray voltage 2300 V, s-lens RF level 60 %, capillary temperature 220 °C, scans 350-1800 m/z. Peptides were separated by 15 cm analytical RSLC column (Acclaim™ PepMap™ 100 C18 2 μm pore size, 150 mm length, 50 μm i.d.) with the gradient of 0-95 % acetonitrile with 0.1 % formic acid): 0-5 % for 5 min, 5-25 % for 40 min, 25-40 % for 10 min, 40-95 % for 2.5 min, and then at 95 % for another 2.5 min. The ten most intense ions from full scan were selected for tandem mass spectrometry. The normalized collision energy was 35 V and default charge state was 2 in HCD mode. Scans were collected in positive polarity mode. Tandem mass spectra were extracted by Mascot Distiller 2.7 (v2.4.1) and all MS/MS samples were analyzed using Mascot (Matrix Science, London, UK; version 2.4.1). Mascot was set up to search against *A. mellifera* and DWV protein databases with the digestion enzyme, trypsin. Mascot was searched with a fragment ion mass tolerance of 0.050 Da and a parent ion tolerance of 10.0 PPM. Carbamidomethyl of cysteine was specified as a fixed modification and oxidation of methionine was specified as a variable modification. Scaffold (version Scaffold_4.9.0, Proteome Software Inc., Portland, OR) was used to validate MS/MS based peptide and protein identification. Peptide identification was accepted if FDR was < 0.01 by the Scaffold Local FDR algorithm. Protein identification was accepted if FDR was < 0.01 and it contained at least two identified peptides. Protein probabilities were assigned by the Protein Prophet algorithm (Nesvizhskii et al., 2003). Proteins containing similar peptides that can not be distinguished based on MS/MS analysis alone were grouped to satisfy the principles of parsimony. Proteins sharing significant peptide evidence were grouped into the clusters.

### Statistical analysis

All experiments were repeated three times except testing the effects of pre-incubation of pupal head cells by purified P-domain protein was repeated four times. All data presented were from representative independent experiments. Statistical analyses were performed with Bell Curve for Excel (Social Survey Research Information Co., Ltd.) and no data point was excluded. The applied statistical tests and P-values are described in figure legends.

## Conflict of interest statement

The authors declare no conflict of interest.

## Author contribution

TK conceived and designed research strategy. YW and JL performed the experiments. YW and TK analyzed data and wrote the paper.

## Funding

This work was supported by Jinji Lake Double Hundred Talents Programme and XJTLU Research Development Fund (RDF-15-01-25) to TK.

## Acknowledgements

We thank Xinyi Li, Ziyun Zhang, Xiangzhen Wei for their contributions to conduct the experiments.

---

**Supplementary Table.**
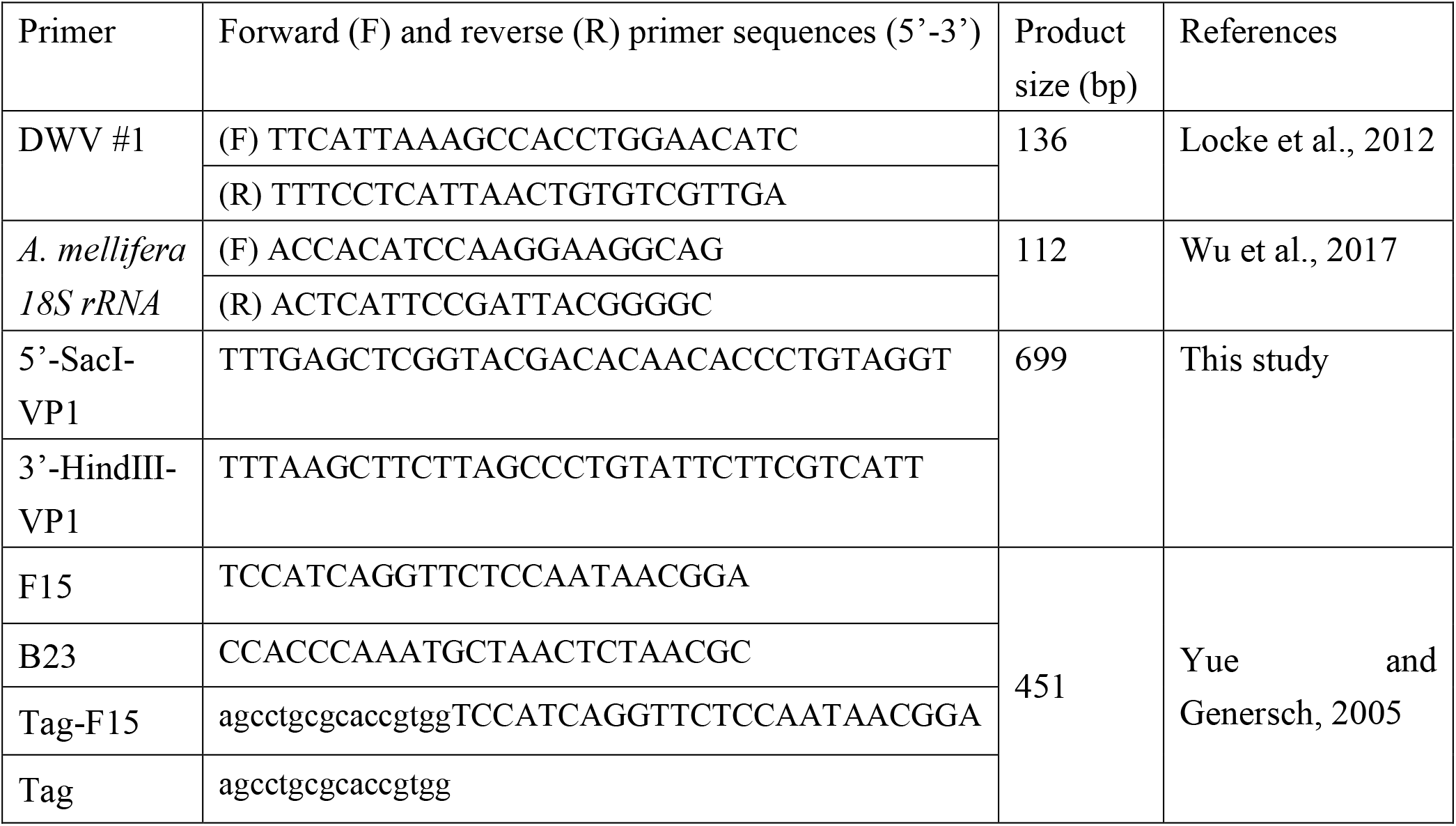
List of primers used in this study.

**Supplementary Figure.**
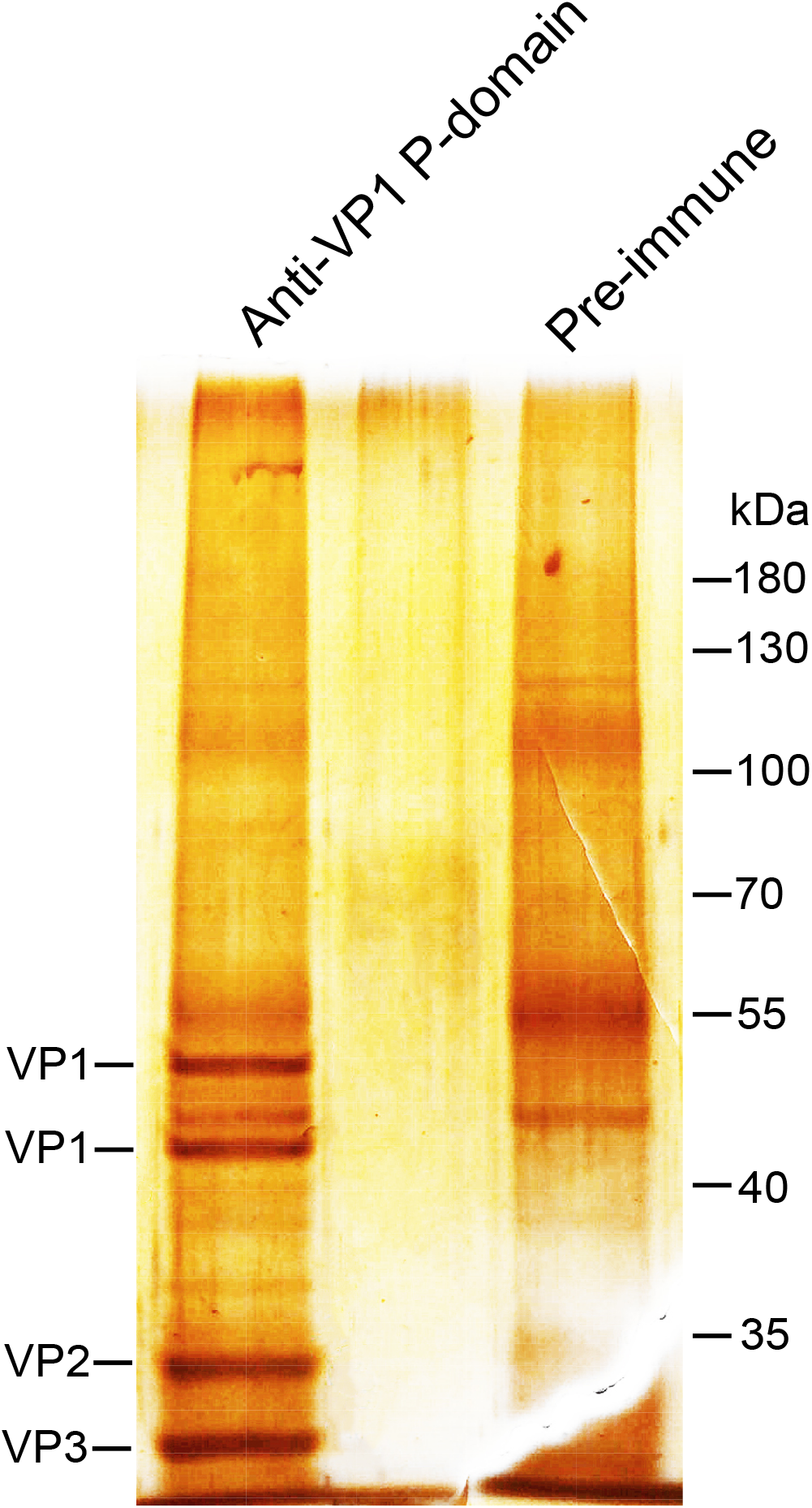
Silver staining of immunoprecipitates by anti-VP1 P-domain antibody. Lysates of DWV-infected pupae were immunoprecipitated by either anti-VP1 P-domain antibody or the pre-immune serum followed by 8 % SDS-PAGE and silver staining. Four major bands specifically immunoprecipitated by anti-VP1 P-domain antibody were identified to be VP1, VP2, and VP3 by the MS analysis.

## Notes

### Competing Interest Statement

The authors have declared no competing interest.

